# SIRT6 as a transcriptional coactivator of GATA4 prevents doxorubicin cardiotoxicity independently of its deacylase activity

**DOI:** 10.1101/725044

**Authors:** Minxian Qian, Linyuan Peng, Zuojun Liu, Xiaolong Tang, Zimei Wang, Baohua Liu

## Abstract

Activity dependent and independent functions for some enzymes are indispensable as significant biological regulators. Deacylase SIRT6 is well-known to improve stress resistance and promote lifespan extension through enzymatic activity-dependent gene silencing. However, whether and how SIRT6 non-enzymatically actives the transcriptional output hasn’t been characterized. Here, we revealed SIRT6 as a coactivator of GATA4, an essential transcription factor for postnatal cardiomyocyte survival, promoting the expression of anti-apoptotic gene. Chemotherapeutic drug, doxorubicin (DOX), remarkably and rapidly decreased SIRT6 expression, leading to transcriptional repression of GATA4 and cardiomyocyte apoptosis. Interestingly, SIRT6 interacted with GATA4 yet enhanced GATA4 acetylation independent of its deacylase activity, by recruiting the acetyltransferase Tip60 to form a trimeric complex. Nonacyl-mimetic mutation of GATA4 thoroughly blocked its ability against DOX cardiotoxicity. Moreover, *Sirt6* transgenic mice exhibited preserved cardiac function with attenuated GATA4 activity in response to DOX. Thus, our studies uncover a previously unrecognized role of SIRT6 in cardioprotection independently of deacylase activity, providing the molecular basis to prevent chemotherapeutic side effects.

## Introduction

Anthracyclines including doxorubicin (DOX) are widely used as the highly efficacious cancer chemotherapeutic agents, but cause severe dose-dependent cardiotoxicity (1,2). Anthracycline-induced heart failure is largely due to myocyte apoptosis (3). GATA4, a member of the GATA transcription factor family with the ability to bind the consensus DNA sequence “GATA” (4), plays important roles in heart development (5,6). As an early-responsive event to DOX, GATA4 is rapidly and dramatically downregulated in transcript and protein levels (3,7–9). GATA4 activates anti-apoptotic gene transcription, such as *Bcl-2* and *Bcl-xL*, against DOX-induced myocyte death (3,10–12). Genetically or pharmaceutically restoring GATA4 has been proposed as an accessible approach to prevent against DOX-induced cardiotoxicity (3,10,13).

SIRT6, a member of a highly conserved sirtuin family of NAD^+^-dependent enzymes, catalyzes histones and a variety of non-histone substrates for deacetylation, thereby controlling chromatin stability and transcriptional restriction (14–16). Through these functions, SIRT6 maintains organismal health and protects against aging and disease pathologies (17–20), such as cancer and metabolic disorders. In particular, SIRT6 has been characterized to prevent the development of cardiac hypertrophy and failure, via deacetylating H3 Lys-9 to repress IGF-Akt signaling (21,22) as well as NF-κB signaling (23,24). Cardiac *SIRT6* expression is sensitive to stress stimuli, like angiotensin II, isoproterenol and ischemia/reperfusion-induced ROS and DOX exposure (22,25–27). Exercise during pregnancy enhances the expression of SIRT6 in neonatal cardiomyocyte against DOX toxicity (28). However, the mechanisms underlying SIRT6 protects against DOX-induced heart failure remain unknown.

Large-scale sequencing of genomes has revealed about 10% of mammalian enzymes have active site-inactivating homologues, however the non-catalytic function has been rather overlooked (29). A human SIRT1 splice variant, which lacks deacetylase coding sequence, binds with p53 and regulates cancer-related gene expression (30). SIRT1 has also been shown to enhance glucocorticoid receptor transcription independently of its deacetylase activity (31). Moreover, catalytically inactive HDAC3 mutants, a class II HDACs, are sufficient to rescue hepatosteatosis in *HDAC3*-deficient mouse liver (32) and aberrant endothelial-to-mesenchymal transition in *HDAC3*-null outflow tracts and semilunar valves (33), through its binding partners. Interestingly, SIRT6 serves as a corepressor of MYC to control cancer cell proliferation, but hardly alters MYC acetylation and H3K9ac on the promoter region of ribosomal protein genes (34). Here, we revealed a novel function of SIRT6 against DOX cardiotoxicity, which acts as a co-activator of GATA4 to upregulate anti-apoptotic gene expression. *Sirt6* transgenic mice preserves cardiac function with attenuated GATA4 activity upon DOX administration.

## Results

### SIRT6 enables GATA4 activity on anti-apoptotic gene expression

Cardiomyocyte apoptosis is the leading cause of heart failure (3). Transcription factor GATA4 is indispensable for postnatal cardiomyocyte survival against DOX (3,10–12), by directly targeting on the promoter regions of anti-apoptotic genes, such as *Bcl-2* and *Bcl-xL*. Likewise, SIRT6 protects cardiomyocytes against hypoxic stress partially by upregulating Bcl-2 proteins by an unknown mechanism (35). Our previous RNA-seq analysis in *Sirt6* wild-type and null MEF cells uncovered a cluster of downregulated genes, including *Bcl-2* and *Bcl-xL* (20) (Supplementary Figure S1A). Cardiac-specific depletion of *Sirt6* in adult mice by a tamoxifen-inducible α-myosin heavy chain (Myh6)-Cre system results in heart failure (22). Quantitative PCR showed *Sirt6* depletion in mouse hearts significantly reduced *Bcl-2* and *Bcl-xL* expression, while neither mRNA nor protein levels of Gata4 were affected (Figure 1A and Supplementary Figure S1B). When we knocked down *Sirt6* or *Gata4* in H9C2 cardiomyocytes by siRNAs, the drastic decline of *Bcl-2* expression was observed, but not *Bcl-xL* (Supplementary Figure S1C). As DOX treatment suppresses anti-apoptotic gene expression by GATA4 depletion (3,7–9), we then performed the time course treatment of DOX in H9C2 cells. Strikingly, *Sirt6* mRNA levels rapidly and dramatically decreased along with the decline of *Gata4*, *Bcl-2*, and *Bcl-xL* expression (Figure 1B). Moreover, depletion of *Sirt6* by siRNAs aggravated DOX-induced decrease of *Bcl-2* and *Bcl-xL* transcripts (Figure 1C). *Sirt6* transgenic mice with the enhanced expression of *Bcl-2* and *Bcl-xL* protected against the decline by DOX induction (Figure 1D). These data implicate that *SIRT6* downregulation by DOX contributes to blocking anti-apoptotic gene expression.

**Figure 1.**
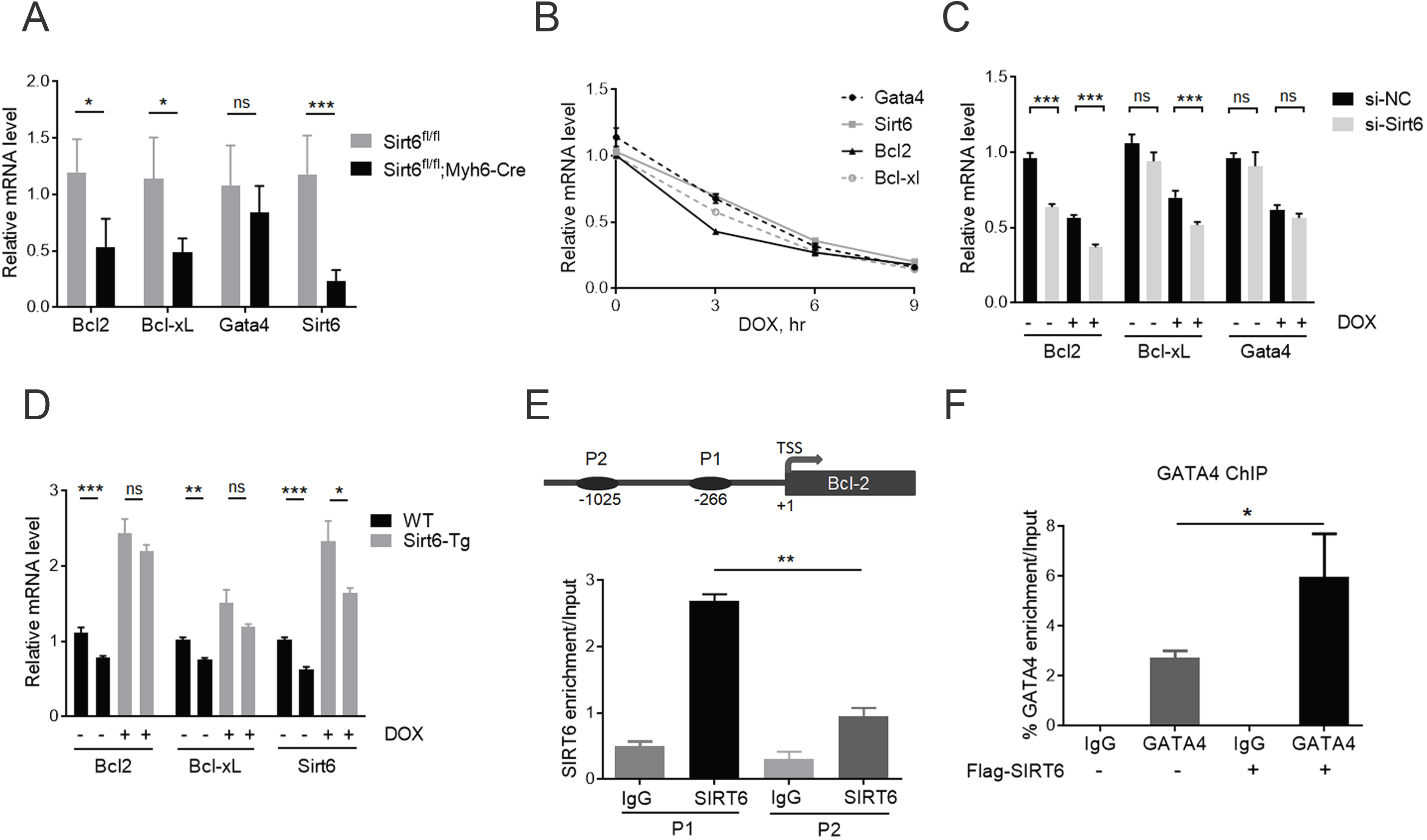

Next, we sought to figure out how SIRT6 cooperates with GATA4 to regulate anti-apoptotic gene expression. GATA4 regulates cardiac *Bcl-2* transcription by binding to the proximal −266 GATA motif (P1) instead of the distal −1025 GATA site (P2) (11) (Figure 1E). We employed a luciferase reporter system cloned with the P1 motif of *Bcl-2* gene. Ectopically expressed SIRT6 significantly elevated GATA4-driven luciferase expression in HEK293 cells (Supplementary Figure S1D). Moreover, chromatin immunoprecipitation (ChIP) demonstrated significantly higher SIRT6 enrichment on P1 locus than on P2 site (Figure 1E), suggesting co-occupancy of GATA4 and SIRT6 on the *Bcl-2* promoter. Indeed, overexpressed SIRT6 enhanced GATA4 occupancy on P1 region (Figure 1F). SIRT6 targets H3K9ac for deacetylation on promoter region to repress gene expression (36). Further ChIP assay using H3K9ac antibodies showed that the acetylation level of H3K9 was merely changed on the P1 region of *Bcl-2* promoter despite that SIRT6 was ectopically expressed (Supplementary Figure S1E). Furthermore, chromatin-fraction analysis showed that overexpressed SIRT6 enhanced GATA4 association with chromatin, which was abrogated in *SIRT6* knockout (KO) HEK293 cells, generated by CRISPR-Cas9 system (Supplementary Figure S1F-G). Together, these results suggest that SIRT6 synergistically enhances GATA4 activity to regulate anti-apoptotic gene expression.

### SIRT6 physically interacts with GATA4

As shown above, SIRT6 and GATA4 shared the P1 promoter region to coordinate *Bcl-2* expression, implicating that two proteins may directly interact. Indeed, GATA4 was detected in the anti HA-SIRT6 immunoprecipitates (IPs) (Figure 2A). Reciprocally, SIRT6 was present in the purified pool of Flag-GATA4 (Figure 2B). The C-terminus of SIRT6 regulates subcellular localization and substrate recognition, while its N-terminus is linked to chromatin association and enzymatic activity (37). Our domain-mapping data showed that the C-terminal deletion of SIRT6 completely abolished its association with GATA4 (Figure 2C). *In vitro* pull-down assay was examined by using purified *E. coli*-expressed His-tagged GATA4 and GST-tagged SIRT6, and the result confirmed the direct interaction of GATA4 with the C-terminus of SIRT6 (Figure 2D). To identify the interaction under physiological conditions, we performed immunofluorescence staining and endogenous IP in H9C2 cells. GATA4 and SIRT6 interacted and were colocalized predominantly in the nucleus (Figure 2E-F). GATA4 possesses two zinc finger domains; C-terminal one is essential for DNA and cofactor binding (38,39) (Figure 2G). Co-immunoprecipitation data showed that SIRT6 specifically bound to the C-terminal zinc finger region (a.a. 251-350) of GATA4 (Figure 2H). Thus SIRT6 might affect DNA binding affinity of GATA4.

**Figure 2.**
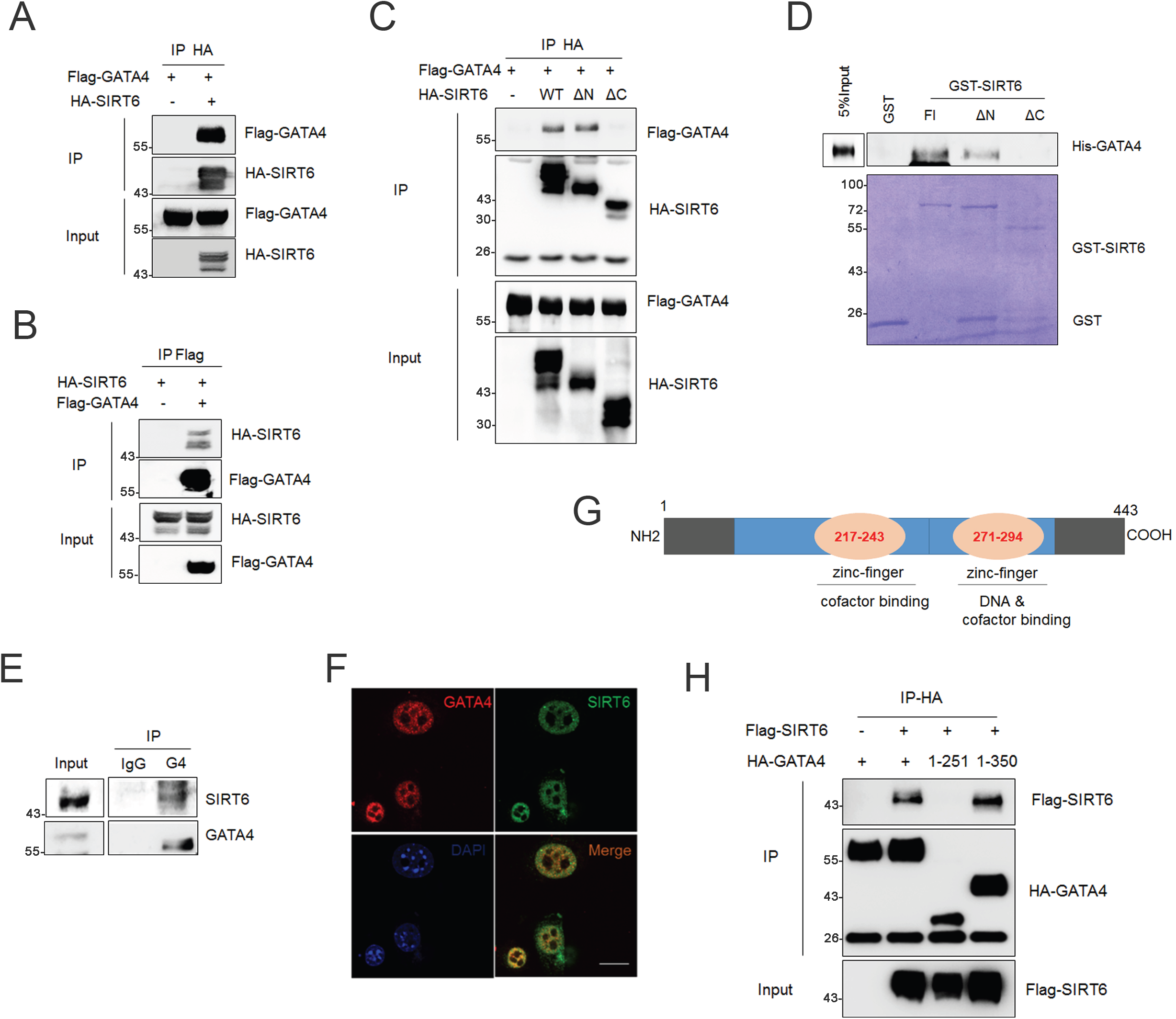

### SIRT6 increases GATA4 acetylation independently of deacylase activity

Posttranslational modifications are critical to the DNA binding capacity and activation of GATA4, especially phosphorylation and acetylation (40–42). Sirtuins (SIRT1-7), as class III histone deacetylases (HDACs), remove the acetyl groups from histones or non-histone proteins. We next addressed whether the acetylation of GATA4 was modulated by the nuclear sirtuins, like SIRT1, SIRT2, SIRT6 and SIRT7. We co-expressed GATA4 with each nuclear sirtuin in HEK293 cells and examined the acetylation level of GATA4 using anti acetyllysine antibody after IP. Unexpectedly, the acetylation level of GATA4 was dramatically increased when SIRT6 was ectopically expressed, while little change was detected in other groups (Figure 3A). Consistently, the lower acetylation level of GATA4 was observed in *SIRT6* depleted cells (Supplementary Figure S2A). Interestingly, the increase of GATA4 acetylation by SIRT6 was counteracted by HDAC2 overexpression, which is shown able to deacetylate GATA4 to modulate embryonic myocyte proliferation (43). (Supplementary Figure S2B). The C-terminal Zn-finger of GATA4 is a key element for DNA and cofactor binding and frequently subjected to acetylation (39). Deletion of the C-terminal Zn-finger domain totally abolished the acetylation and chromatin binding ability of GATA4, as determined by anti-AcK and anti-histone H3 immunoblotting in the GATA4 IPs (Figure 3B). Notably, ectopically expressed SIRT6 increased GATA4 acetylation in this region (Figure 3B), explaining the way how SIRT6 regulates DNA binding activity of GATA4.

**Figure 3.**
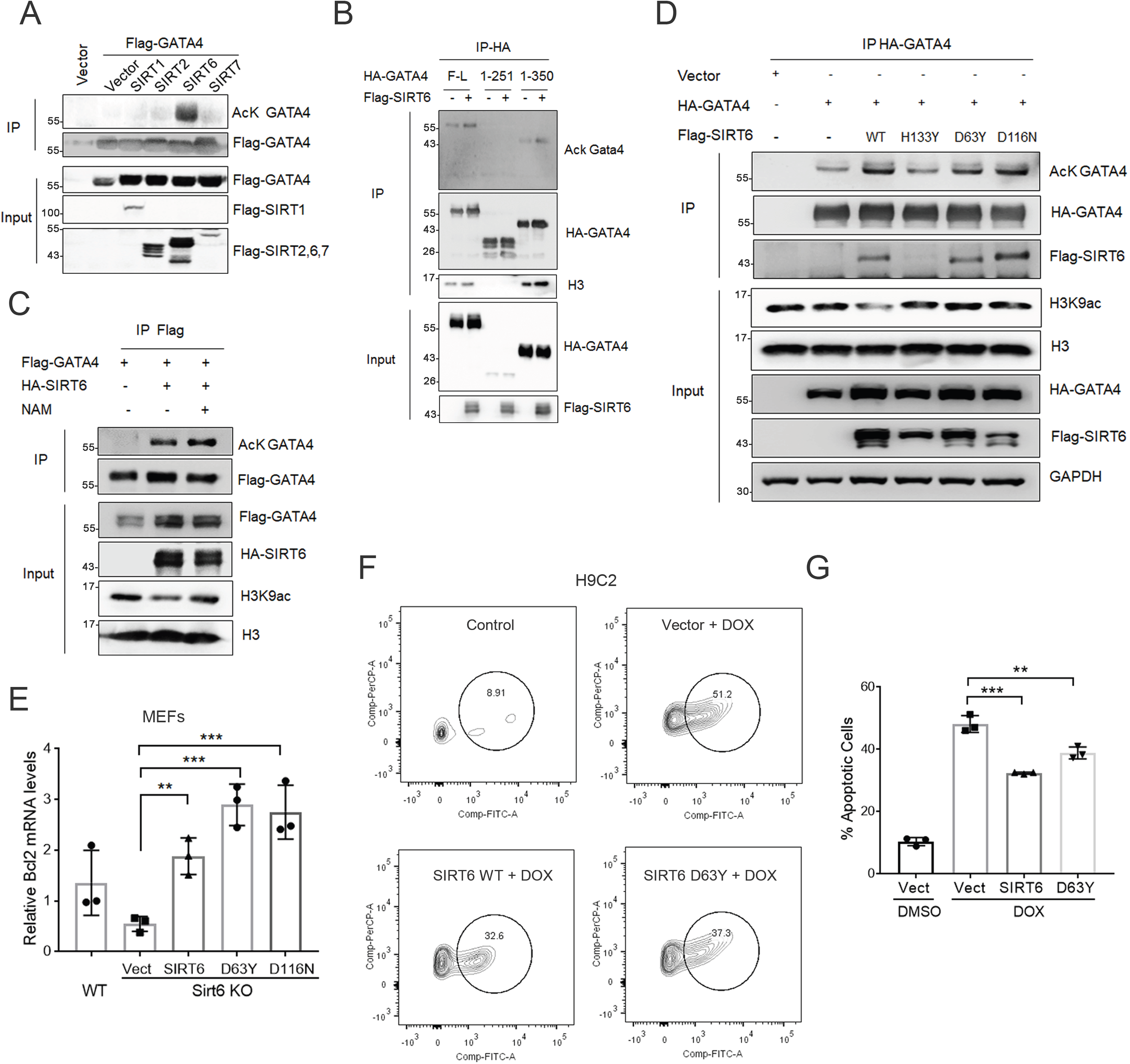

SIRT6 possesses not only deacetylase activity, but also defatty-acylase and mono-ADP-ribosyl transferase activities. We thus asked whether the increase of GATA4 acetylation requires SIRT6 enzymatic activity. We utilized nicotinamide (NAM) to block deacylase and mono-ADP-ribosylase activities (44,45), and then investigated the effect on GATA4 acetylation. As shown, the NAM treatment abrogated H3K9 deacetylation mediated by sirtuins, but was unable to inhibit the increased acetylation level of GATA4 induced by SIRT6 overexpression (Figure 3C). More importantly, the overexpression of catalytically inactive mutations of SIRT6, including D63Y and D116N (25), were still capable to bind and enhance GATA4 acetylation (Figure 3D). By contrast, SIRT6 H133Y mutation failed to increase GATA4 acetylation, most likely due to the extinct binding affinity of H133Y SIRT6 to GATA4 (Figure 3D and Supplementary Figure S2C). Moreover, reintroduction of SIRT6 wild-type, D63Y or D116N mutation largely rescued the expression of *Bcl-2* and *Bcl-xL* in *Sirt6* null MEF cells (Figure 3E and Supplementary Figure S2D), and attenuated DOX-induced apoptosis in H9C2 cells by flow cytometry analysis (Figure 3F and Supplementary Figure S2E). The data strongly suggest that SIRT6-enhanced transcription activity of GATA4 is independent of its catalytic activity.

### SIRT6 recruits Tip60 for GATA4 acetylation

The finding that SIRT6 interacts with GATA4 and enhances its acetylation apparently indicates that additional factors are involved. To figure out which acetyltransferase is responsible for it, we examined the alteration on SIRT6-regualted GATA4 acetylation under treatment with various HAT inhibitors, including Cpth2 for Gcn5 (46), Mg149 for Tip60 (47) and C646 for p300 (48,49). Interestingly, Mg149 treatment remarkably blocked the increase of acetylation of GATA4 induced by SIRT6, while both Cpth2 and C646 failed (Figure 4A). While p300 catalyzes GATA4 acetylation during hypertrophic stimuli in cardiomyocytes (30,43), Tip60 is most likely involved in SIRT6-enhanced GATA4 acetylation. Specific deletion of *Tip60* in heart results in a shortened lifespan, accompanied with the increased myocyte density and apoptosis (50). We next sought to evaluate whether GATA4 is a target of Tip60. As shown, ectopically expressed Tip60 increased the acetylation level of GATA4, which was abolished by the inhibitor Mg149 (Supplementary Figure S3A). Knocking down *Tip60* by siRNAs remarkably decreased the acetylation level of GATA4 (Supplementary Figure S3B). To confirm the function of Tip60 in SIRT6-enhanced GATA4 acetylation, we knocked down *Tip60* in HEK293 cells. As shown, the increase of GATA4 acetylation induced by SIRT6 was significantly dampened (Figure 4B). Interestingly, Tip60 overexpression was inefficient to acetylate GATA4 in *SIRT6*-depleted HEK293 cells (Figure 4C).

**Figure 4.**
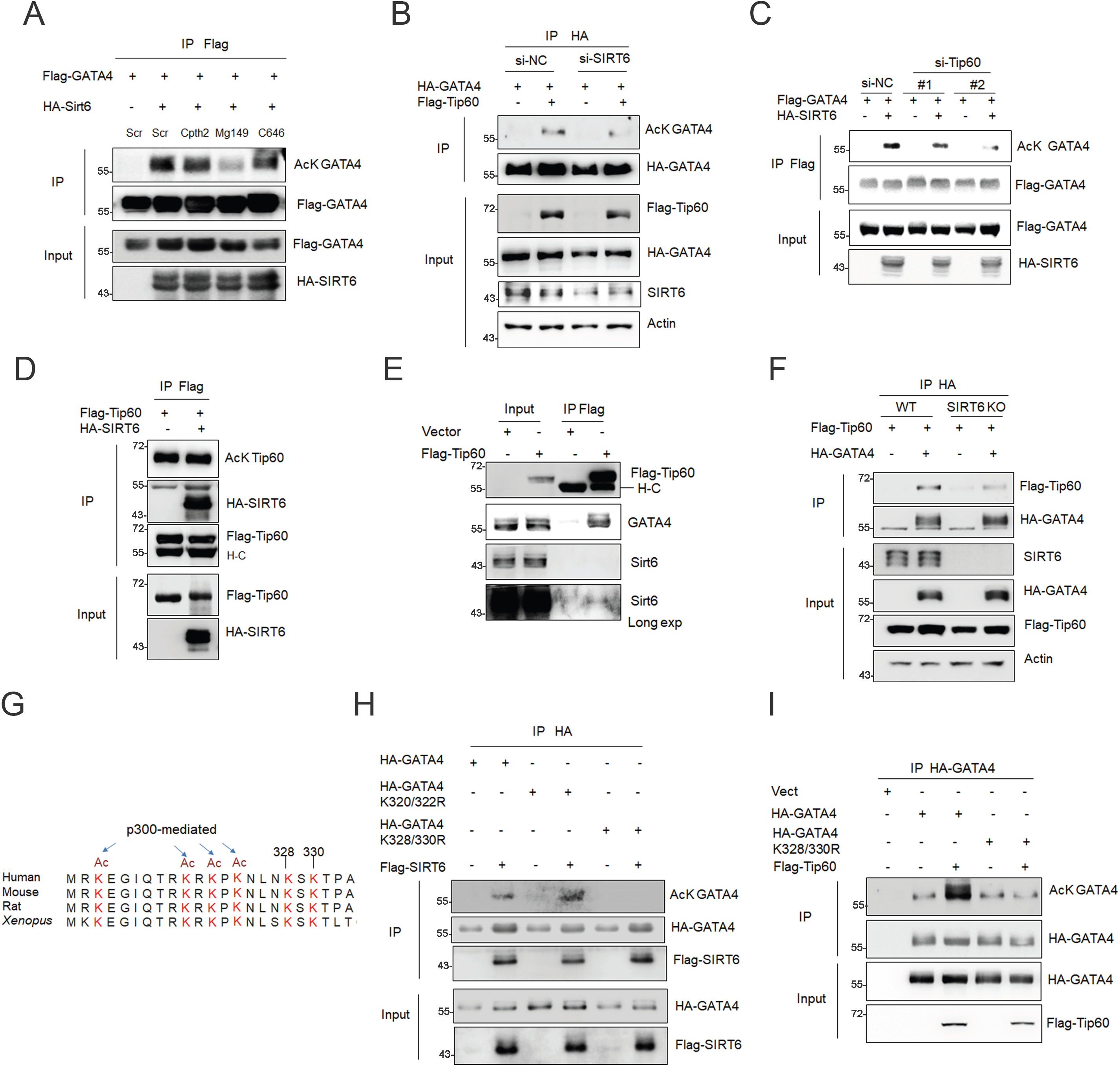

SIRT6 deacetylates Gnc5 to activate its acetyltransferase activity, which then acetylates PGC1α (51). Autoacetylation activates Tip60, which is negatively regulated by SIRT1 (52). As shown, overexpressed SIRT6 hardly affected acetylation level of Tip60 (Figure 4D), although SIRT6 and its enzyme-dead mutations could bind with Tip60 identified by Co-IP (Supplementary Figure S3C). Both SIRT6 and GATA4 were present in the anti Flag-Tip60 immunoprecipitates (Figure 4E). However, the interaction between Tip60 and GATA4 was undetectable in *SIRT6* KO HEK293 cells (Figure 4F). The data further support that SIRT6 regulates Tip60-mediated GATA4 acetylation in a deacetylase-independent manner.

As demonstrated above, SIRT6 bound and enhanced the acetylation of GATA4 on C-terminal Zn-finger domain, which is highly and multiply modified (53,54). Within this region, there are six evolutionally conserved lysine (K) residues (Figure 4G). Of these Ks, p300 acetylates GATA4 on K313/320/322/324 and thus enhances the DNA-binding activity, resulting in stress-induced cardiac hypertrophy (39,55). By K-to-R mutation, we identified K328/330 mutant (2KR), which almost abolished the increase of GATA4 acetylation by overexpression of SIRT6 or Tip60 (Figure 4H-I). The data indicate that SIRT6/Tip60 mediates acetylation of GATA4 on K328/330.

### Acetylation is critical for GATA4 activity and protection against DOX

Restoring GATA4 transcription activity has been recognized as a promising approach to prevent DOX-induced heart failure (9,10). Since SIRT6 bound and enhanced GATA4 acetylation, we next investigated the cardioprotective roles of SIRT6/Tip60-mediated GATA4 acetylation. We first co-expressed HA-GATA4 and Flag-SIRT6 in HEK293 cells, and then treated the cells with DOX. HA-GATA4 was purified by IP and the levels of Flag-SIRT6 protein and GATA4 acetylation were tested by Western blotting. We found that DOX disrupted the interaction of GATA4 and SIRT6, and simultaneously reduced the acetylation level of GATA4 in a dose-dependent manner (Figure 5A). We next generated H9C2 cell lines stably expressing wild-type (WT), K328/330R (2KR)-or K328/330Q (2KQ)-mutated forms of GATA4 (Supplementary Figure S4A). Over-expressed GATA4 attenuated the decline of *Bcl-2* in H9C2 cells subjected to DOX, but the 2KR mutation failed (Figure 5B). Both ChIP and chromatin-binding fraction assay showed that 2KR mutation reduced the DNA-binding capacity of GATA4 (Figure 5C and Supplementary Figure S4B). It was previously reported that overexpressed GATA4 alleviates DOX-induced atrophic reaction and apoptosis in cardiomyocytes (3). This benefit was abrogated by 2KR mutation (Figure 5D-F and Supplementary Figure S4C). Further, colony formation assay demonstrated that GATA4 KR failed to rescue myocyte survival in response to DOX (Figure 5G). These data suggest that acetylation mediated by SIRT6/Tip60 is critical for the cardiac protective role of GATA4.

**Figure 5.**
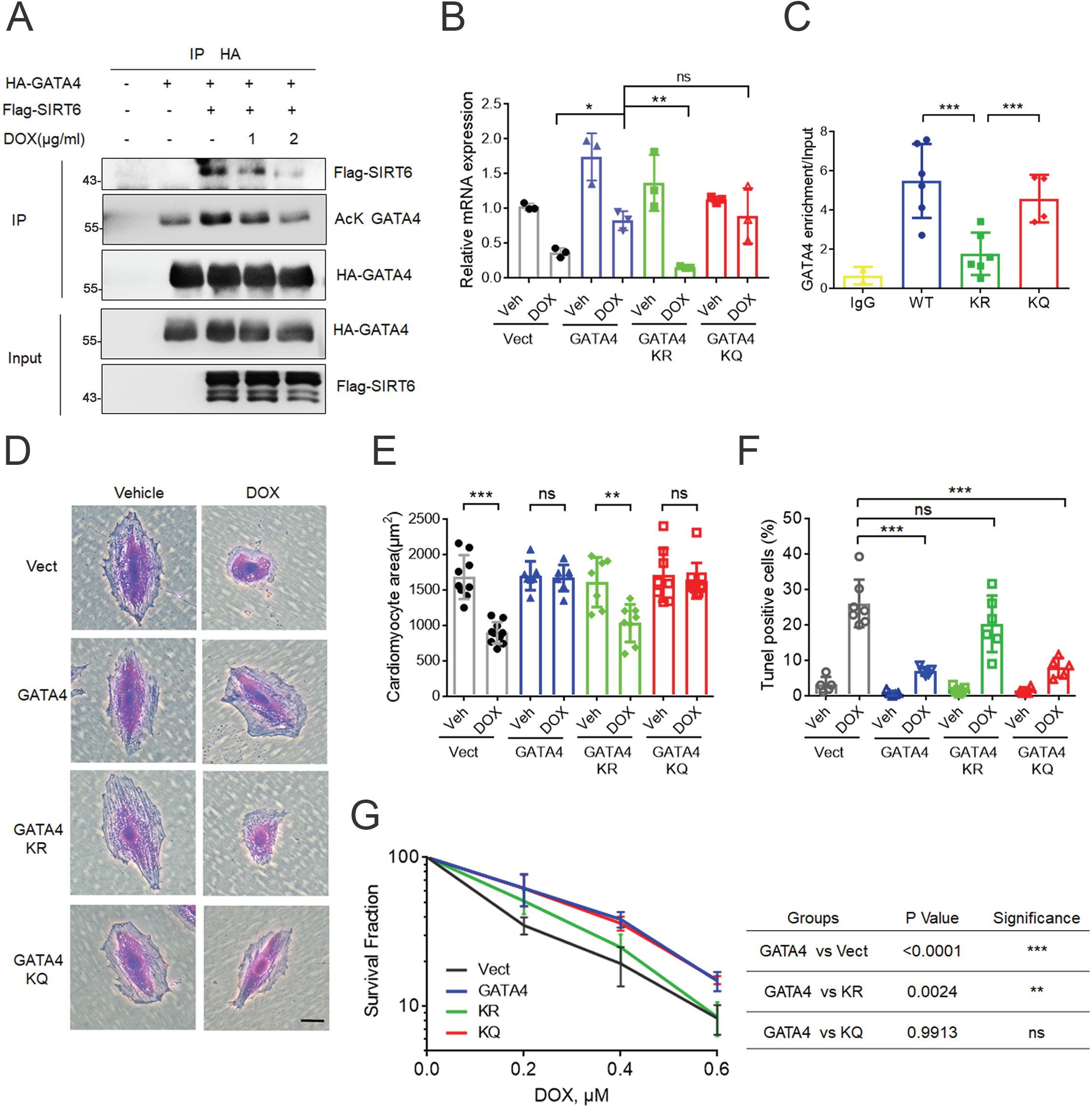

### *Sirt6* transgene alleviates DOX-induced cardiomyopathy

We next examined the function of SIRT6 in preventing DOX cardiotoxicity *in vivo*. Since cardiac-specific or whole-body *Sirt6* deficiency in mice spontaneously develop heart failure (18,22), we here employed *Sirt6* transgenic mice for DOX treatment. WT and *Sirt6* transgenic mice were subjected to acute DOX treatment (a cumulative dose of of 20 mg/kg DOX via intraperitoneal injection twice in one week). Of note, *Sirt6* transgenic mice exhibited well tolerance to DOX-induced mobility (Figure 6A). DOX administration causes body weight loss in cancer patients (56,57). In laboratory mice, *Sirt6* transgene significantly attenuated DOX-induced body weight loss (Figure 6B), and the reduction of heart weight/body weight ratio (HW/BW) and cross-sectional area of cardiomyocytes (Figure 6C-D and Supplementary Figure S5A). Immunohistochemistry staining showed a significant restoration of Gata4 protein in hearts from *Sirt6* transgenic mice after DOX treatment (Figure 6E and Supplementary Figure S5B). In line with the function of GATA4, SIRT6 mitigated DOX-induced cardiac atrophy. The reduction of cardiomyocyte size and the increased fibrosis were largely rescued in *Sirt6* transgenic mice (Figure 6F-H and Supplementary Figure S5C). Consistent with the enhanced expression of *Bcl-2* and *Bcl-xL* by overexpressed Sirt6 (Figure 1D), the number of TUNEL-positive cells in *Sirt6*-transgenic heart (~10.6%) was markedly lower than that in WT (~30.8%) after DOX treatment (Figure 6I and Supplementary Figure S5D). These data support the notion that overexpression of SIRT6 *in vivo* alleviates the extent of DOX-induced acute heart failure and increases survival.

**Figure 6.**
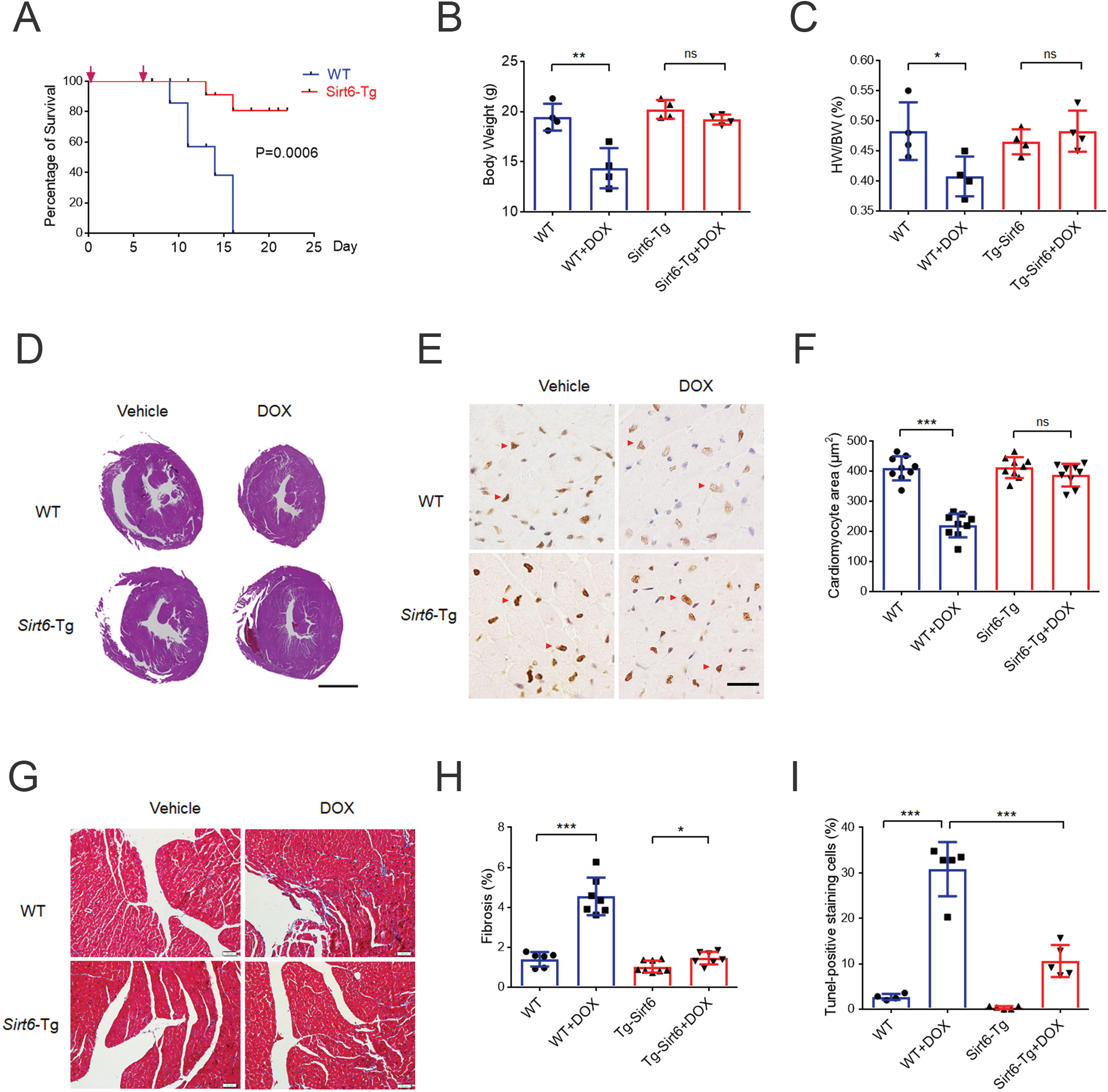

Overall, we showed that SIRT6 prevents the cardiac side effects of DOX by restoring GATA4 activation in a deacylase independent manner. SIRT6 recruits Tip60 to acetylate GATA4, enhancing anti-apoptotic gene expression; upon DOX exposure, the reduction of SIRT6 expression suppresses the chromatin-binding affinity and transcription activity of GATA4, subsequently triggering cardiomyocyte apoptosis (Figure 7).

**Figure 7.**
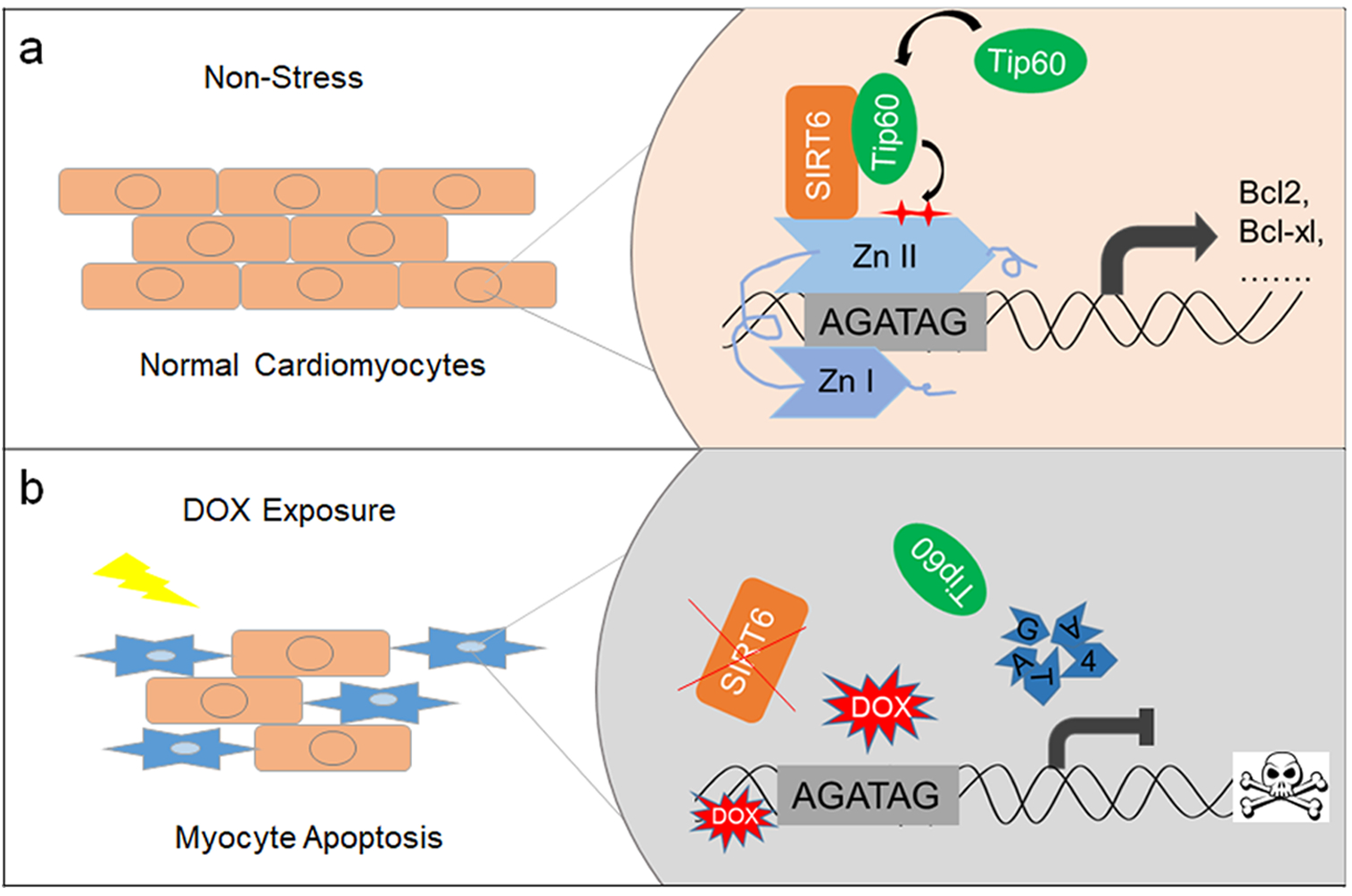

## Discussion

The activity of the silence protein Sir2, as a NAD^+^-dependent histone deacetylase, is highly conserved ranging from yeasts to humans. SIRT6, one member of mammalian sirtuin family, has been well characterized in regulation of metabolism, inflammation, genomic stability, cancer and aging. SIRT6 represses gene expression predominantly through three criteria: 1) catalyzing histone deacetylation to remodel the chromatin structure; 2) altering post-translational modifications and activities of transcription factors; 3) regulating the expression of transcription factors. However, the mechanism of SIRT6-mediated transcriptional activation is still unknown. Most recently, SIRT6 mono-ADP-ribosylation activity was addressed critical for NRF2 target gene activation upon oxidative stress (58). In this study, we for the first time identify a catalytic activity independent role of SIRT6, acting as a coactivator of GATA4 and positively regulating anti-apoptotic gene expression.

In addition to deacetylation, SIRT6 possesses the lysine defatty-acylase and adenosine diphosphate (ADP)-ribosyltransferase activities; all require NAD^+^ as a coenzyme or a substrate (59,60). NAM, a most potent inhibitor of Sir2 by binding to a conserved pocket adjacent to NAD^+^ (61), was insufficient to block SIRT6-enhanced acetylation of GATA4, but H133Y mutation of SIRT6 did. Notably, H133Y not only blocks the activities of SIRT6 but also disrupts its interactions, like histones (37), Ku80 (62) and GATA4 in this study. To strengthen the conclusion, we employed other point mutations, D63Y and D116N, which disrupt the affinity of SIRT6 for NAD^+^ (63). Deacetylase-independent role of SIRT6 in regulating GATA4 was further confirmed. Moreover, we showed that SIRT6 and these mutants recruit Tip60, with no change on Tip60 acetylation, to increase acetylation of GATA4. SIRT6 mutation on Asp63 leads to human perinatal lethality, suggesting an essential role of SIRT6 deacetylase activity in embryogenesis (25), we herein revealed a deacetylase-independent role of SIRT6 ensuring postnatal cardiomyocyte survival.

Tip60 (Tat-interactive protein 60), a member of the MYST family of acetyltransferases, has a crucial function in DNA damage response, apoptosis, and cell cycle control. The acetylation status of Tip60 is intimately linked to its activity (52,64); SIRT1 negatively regulates it (52). Heterozygous *Tip60* mice reproduce normally, whereas homozygous global ablation causes completely penetrant embryo lethality (65). Tip60 is highly expressed at early stages of heart development (66). Heart-specific depletion of *Tip60* in mice increases myocyte density and apoptosis, accompanied by a shortened lifespan (50). However, the underlying mechanism has not been well elucidated. GATA4 is essential for heart development and cardiomyocyte survival. So far, only p300/CBP has been reported to target GATA4 for acetylation, subsequently inducing cardiac hypertrophy (39,55). Here, we found that Tip60 serves as a novel acetyltransferase for GATA4 acetylation on K328/330 to protect against stress-induced heart failure, providing an explanation by which Tip60 maintains heart homeostasis.

The clinical use of DOX, one of the most effective chemotherapeutic agents against a broad spectrum of malignancies, is limited due to its severe side effect cardiotoxicity. Multiple mechanisms are involved in DOX-induced heart failure, including oxidative stress, DNA double-strand breaks and apoptosis (67,68). In particular, GATA4 is shown to be downregulated and fail to protect cardiomyocyte survival in response to DOX (3,10). How DOX triggers GATA4 proteolysis has not been well established. Here we showed that SIRT6/Tip60-mediated acetylation of GATA4 is required for GATA4 DNA binding and transactivation against DOX-induced myocyte apoptosis. *Sirt6* transgenic mice are more resistant to DOX-induced cardiomyopathy. These findings suggest that SIRT6/Tip60-mediated GATA4 acetylation can be considered as a potential target in preventing DOX cardiotoxicity.

