## Supplementary figures and images for "SIRT6 as a transcriptional coactivator of GATA4 prevents doxorubicin cardiotoxicity independently of its deacylase activity"

### Supplemental figures

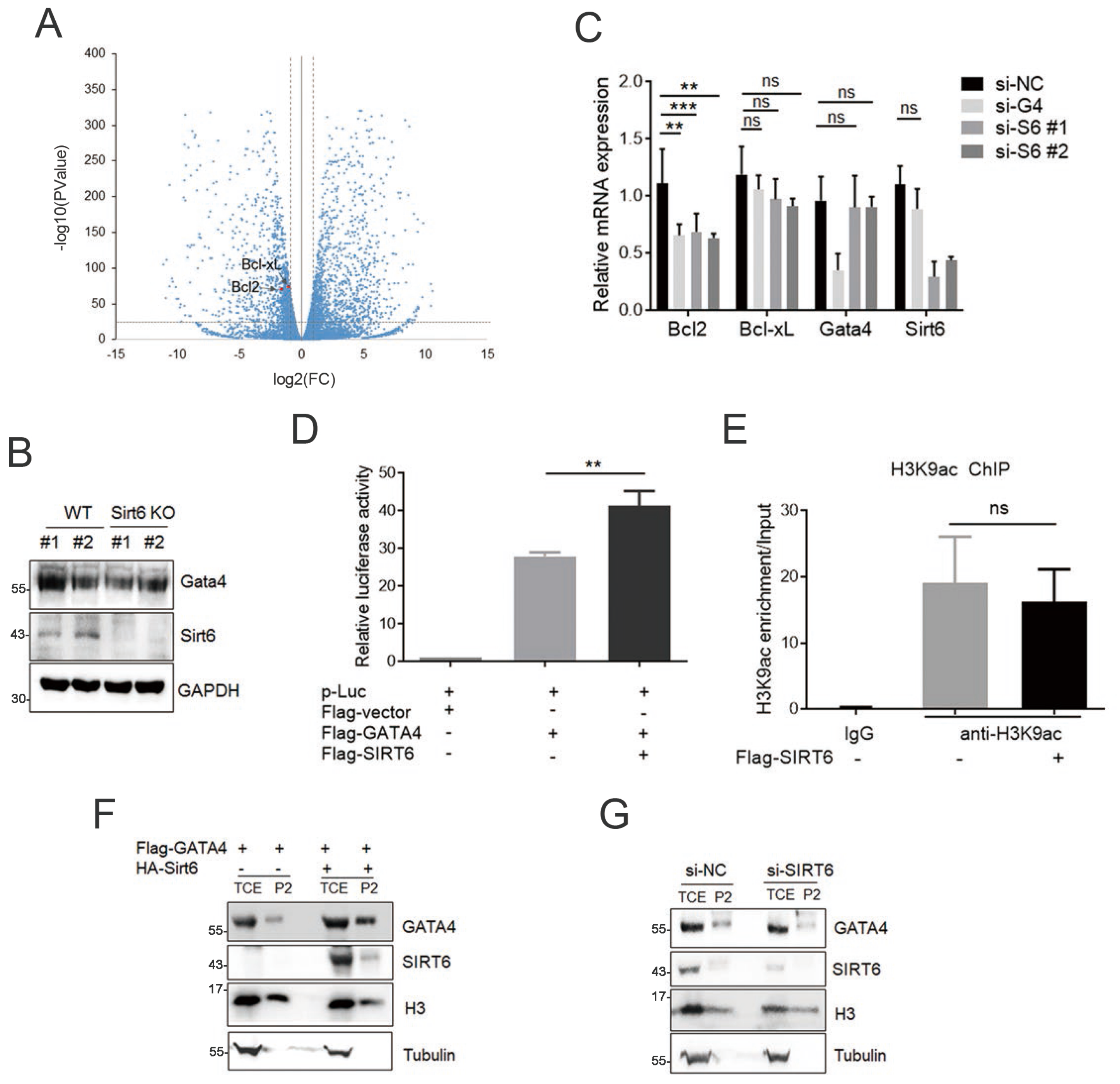

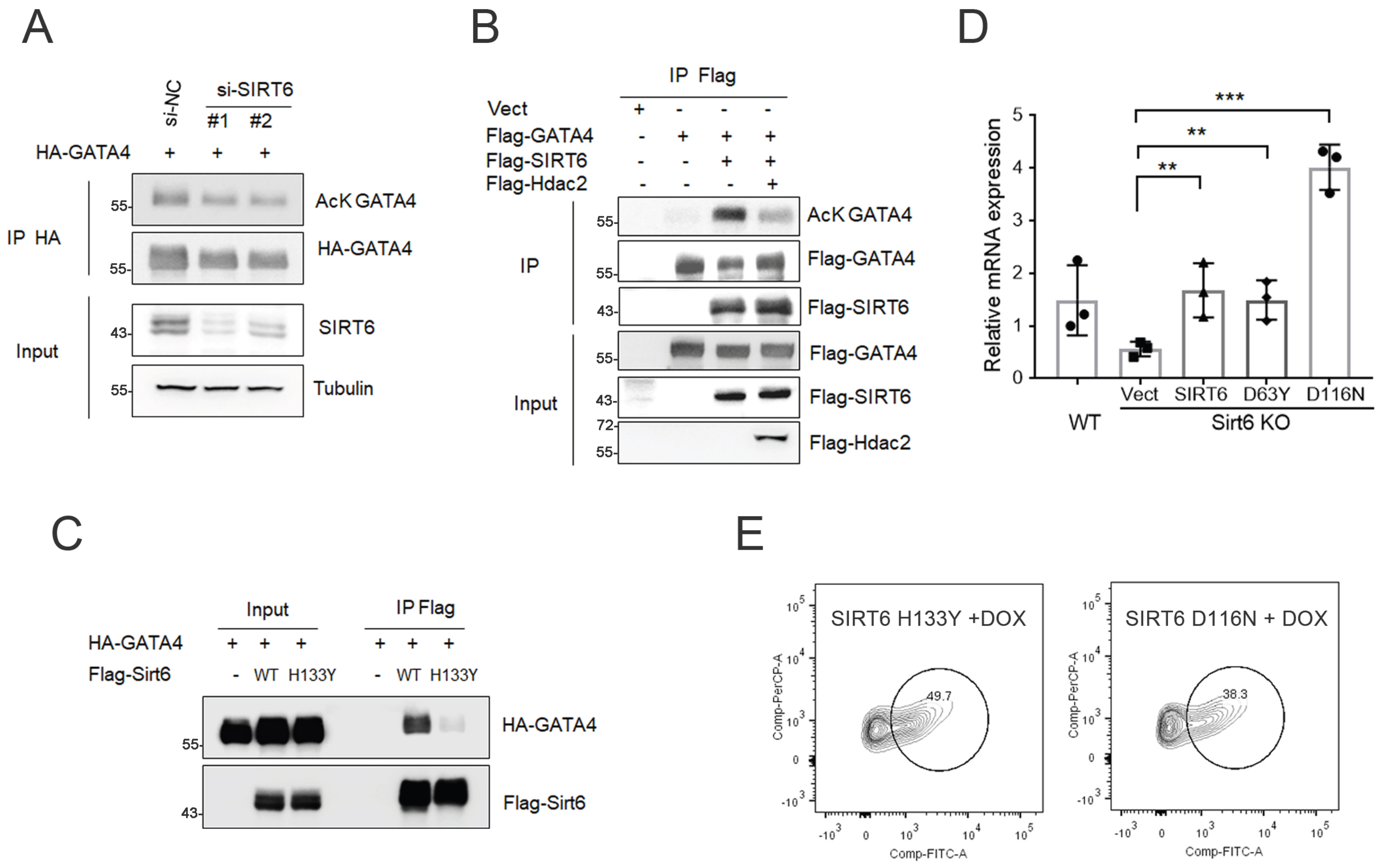

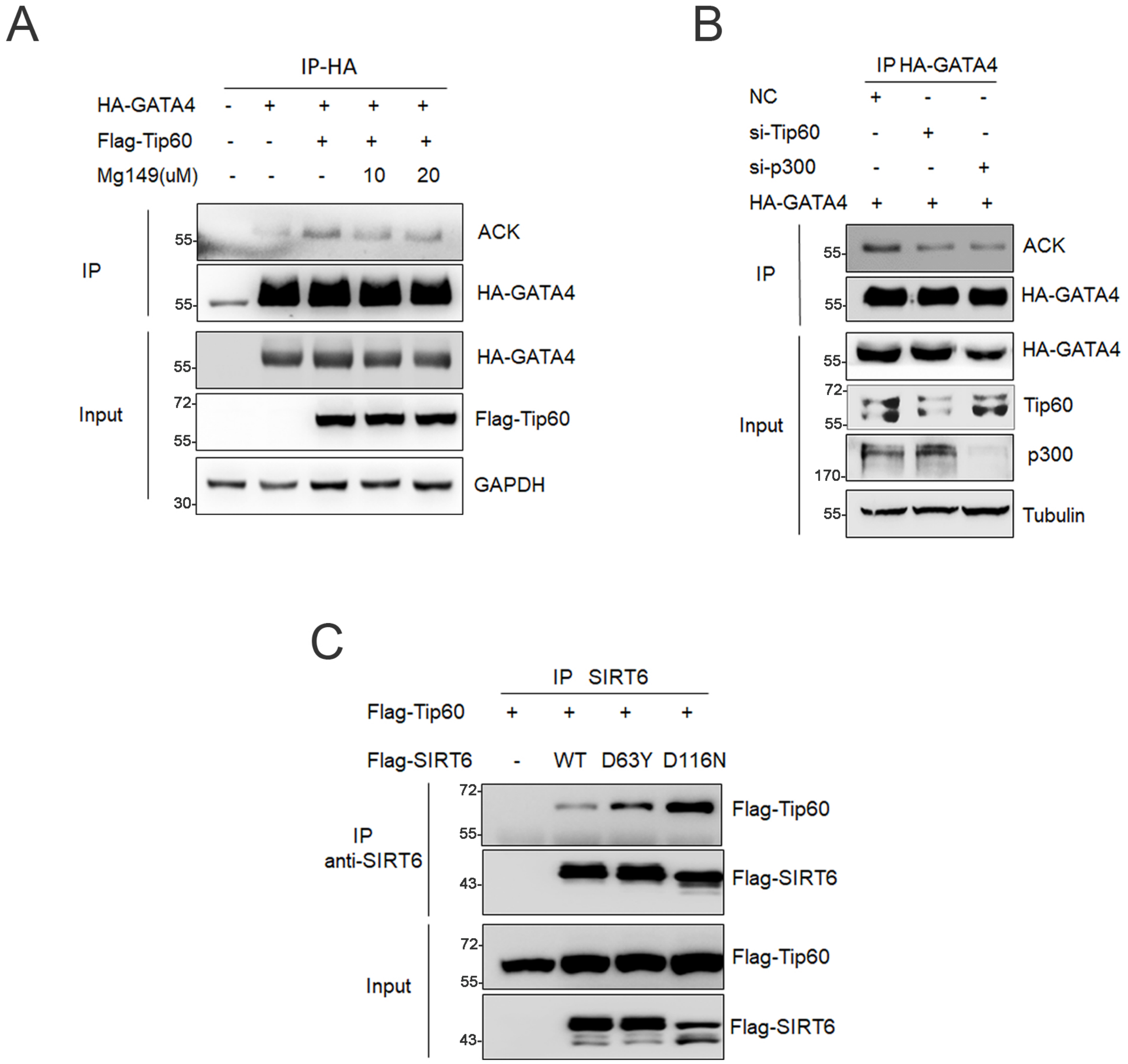

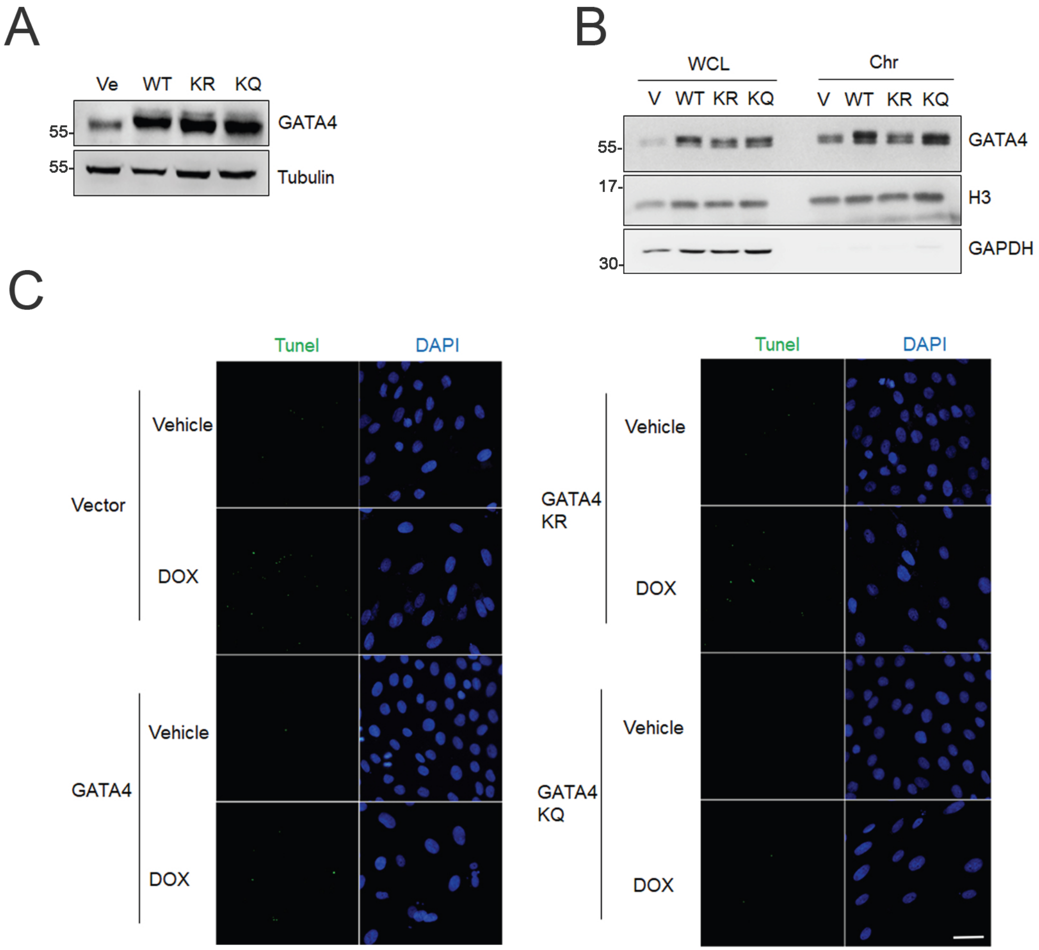

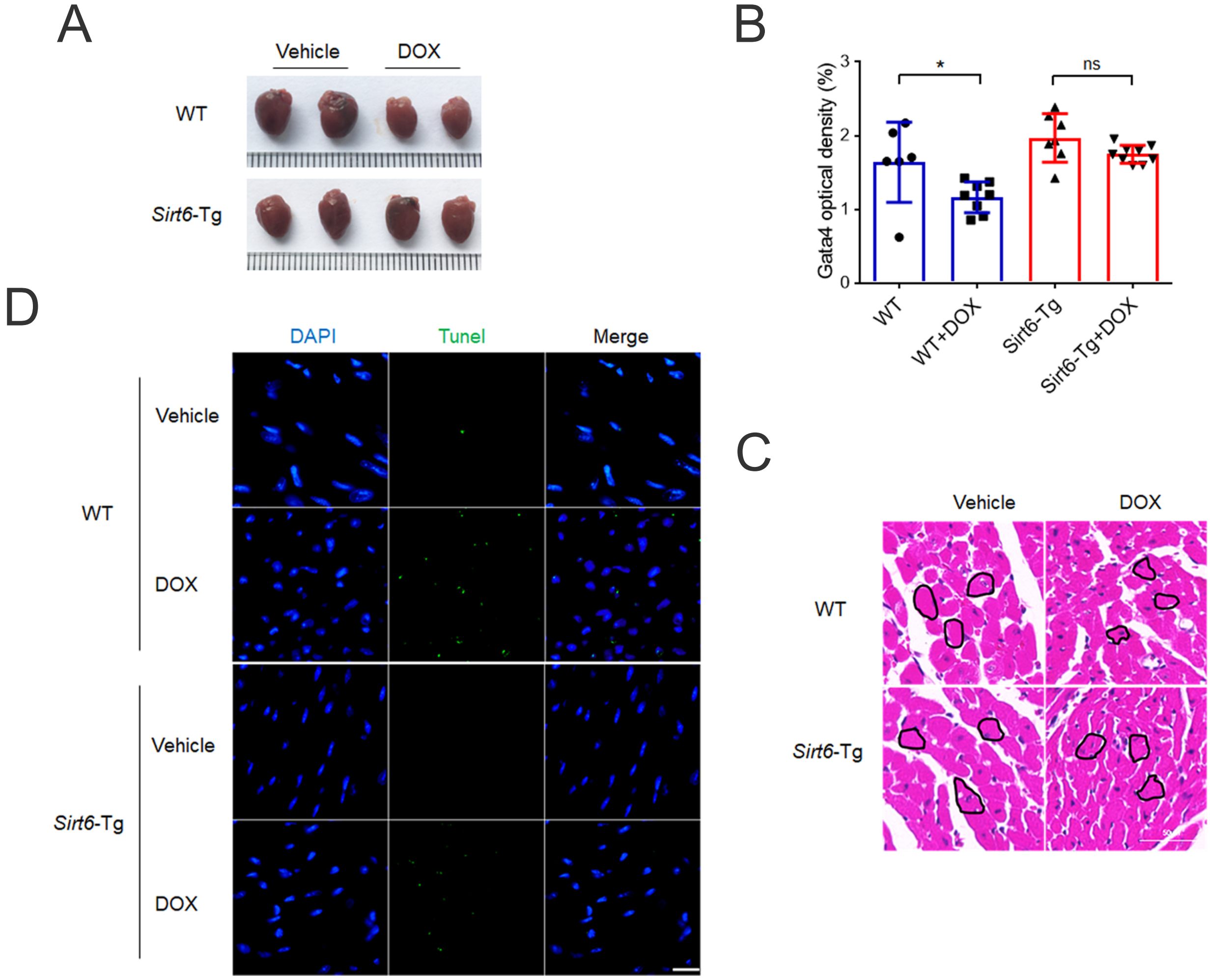
