## Supplementary material for "SIRT6 as a transcriptional coactivator of GATA4 prevents doxorubicin cardiotoxicity independently of its deacylase activity": Materials and methods

#### Cell lines and treatments

HEK293 (CRL-1573) and H9C2 (GNR-5) cells were purchased from ATCC. Wild-type and *Sirt6-/-* MEFs were obtained from the animal embryos. *SIRT6* knockout (KO) HEK293 cell lines were generated using the CRISPR/Cas9 system as described previously (1). All these cell lines were cultured in Dulbecco's modified Eagle's medium (DMEM, Life Technologies, USA) supplemented with 15% fetal bovine serum (FBS), 100 U/ml penicillin and streptomycin (P/S) at 37°C in 5% $CO_2$ and atmospheric oxygen condition. Basically, cells were treated with doxorubicin at the indicated dosages for the specific analysis.

#### Mice and DOX administration

To generate cardiac-specific *Sirt6* deletion mice, $Sirt6^{flox/flox}$ mice were crossed with mice carrying *Myh6-cre/Esr1* transgene. 4-hydroxytamoxifen (4-OHT) was injected intraperitoneally (i.p.) daily for 3 consecutive days. One month later, the heart tissues were harvested and analyzed by PCR-based genotyping and quantitative PCR. *Sirt6*-transgenic mice (*Sirt6*-Tg) has been described previously (1). To test the cardioprotective function of SIRT6 against doxorubicin (DOX), wild-type (WT) and *Sirt6*-Tg mice received intraperitoneal injection of 10 mg/kg DOX twice in one week to induce the chronic cardiotoxicity. Survival and body weight were monitored regularly. Saline treatment was performed as a parallel control. The heart tissues were harvested for histopathological examination two weeks later after the last injection. Mice were housed and handled in the laboratory animal research center of Shenzhen University. All experiments were performed in accordance with the guidelines of the Institutional Animal Care and Use Committee (IACUC).

#### Cell line generation and colony formation assay

To establish stably transfected H9C2 cells, the plasmids of pcDNA3.1, GATA4, GATA4 KR, GATA4 KQ were digested with *PvuI* nuclease for 30min to linearize, and then individually transfected into H9C2 cells with lipofectamine 3000 (Invitrogen L3000015). The medium was changed after 24 hours, and supplemented with 2 mg/ml of G418 for sorting. After 10-day selection, the stably transfected cell lines were obtained and confirmed by western blotting. For colony-formation assay, cells were seeded in 6-well plates in triplicate and cultured under normal growth conditions in the presence or absence of doxorubicin at the indicated doses. After further cultured for 10-14 days, cell colonies were stained the with 0.5% crystal violet solution. The number of colonies in each well was quantitated and the surviving fraction was calculated.

**Plasmids and RNA interference**

Human Flag-GATA4 and Flag-Tip60 were all purchased from Vigene Biosciences. Human Flag-SIRT1, 2, 6, 7 were purchased from Addgene. Flag-GATA4 and Flag-SIRT6 with amino acid substitution mutations were generated by PCR-based mutagenesis using a KOD-PLUS kit (Toboyo). The truncated expression plasmid of SIRT6 or GATA4 was constructed by PCR-based deletion based on the HA-SIRT6 or HA-GATA4 plasmid as a template. The primers used for mutagenesis are listed (see Supplementary Table S1). To downregulate the target gene expression, cells were transfected with small interfering RNAs (siRNAs) for 48 hours using lipofectamine 3000 according to the manufacturer's instructions. A scrambled siRNA was used as a control. All siRNAs were purchased from Gene Pharma, the specific sequences are included in Supplementary Table S1.

**Chromatin bound fraction assay**

The cells were carefully scraped off with 1 ml cold PBS buffer, centrifuged at 3000rpm for 1 min. Re-suspend the cell pellets with 500 μl Buffer A (10 mM HEPES, 10 mM KCl, 1.5 mM $MgCl_2$, 0.34 M Sucrose, 10% Glycerol, 1 mM DTT, 0.1% Triton X-100, 10 mM $Na_3VO_4$, protease inhibitor cock-tail) and keep for 10 minutes on ice. Lysate was centrifuged at 1300 ×g for 5 min. After centrifugation, the supernatant contains cytosolic proteins. The precipitate was washed once with Buffer A, and then lysed in 250 μl Buffer B (3 mM EDTA, 0.2 mM EGTA, 1 mM DTT, 10 mM $Na_3VO_4$, and protease inhibitor cock-tail). After incubation and centrifugation (1700 ×g, 5 min), the pellet (chromatin bound fraction) was obtained and then denatured by 1×SDS loading buffer.

**Immunoprecipitation (IP) and pull-down assay**

Co-immunoprecipitation assay was performed as described previously (2). Briefly, cells were washed with cold PBS and harvested. Suspend the cell pellets with 500 μl of IP lysis buffer (20 mM Tris-HCl PH 8.0, 250 mM NaCl, 5 M sodium pyrophosphate, 1 mM sodium fluoride, 10% glycerol, 2 mM EDTA, 10 mM $Na_3VO_4$, protease inhibitor cocktail, and 1 mM nicotinamide). After sonication and centrifugation, the supernatant was collected and incubated with anti-HA or anti-Flag M2 agarose (Sigma) overnight at 4℃ on a rotating platform. The beads were then washed for three times with 1 mL cold lysis buffer. To denature, 1×SDS loading buffer was added and boiled at 98°C for 6 min. The samples including input (5%) were applied for western blotting.

For *in vitro* pull-down experiment, GST or GST-SIRT6 (1 μg) conjugated GSH beads were suspended in NETN buffer (20 mM Tris–HCl pH 8, 100 mM NaCl, 1 mM EDTA, 0.5% NP40 and phosphatase and protease inhibitor cocktail). 0.2μg purified His-GATA4 proteins were added to GST beads and incubated overnight at 4°C, followed by 3× washes with NETN buffer. Boiled eluates were separated and detected by coomassie staining and immunoblotting.

**Protein extraction and Western Blotting**

Mouse tissues were homogenized in 1 ml of ice-cold tissue lysis buffer (25 mM TrisHCl, pH 7.5, 10 mM $Na_3VO_4$, 100 mM NaF, 50 mM $Na_4P_2O_7$, 5 mM EGTA, 5 mM EDTA, 0.5% SDS, 1% NP-40, and protease inhibitor cock-tail). After sonication, lysates were centrifuged at 16,000 ×g for 10min. The clean supernatant was carefully collected. Protein concentration was determined by bicinchoninic acid kit (BCA, Thermo). Western Blotting were performed following the standard protocol. Anti- acetylated-Lysine (CST, Ac-K-100, #6952), anti-Tip60 (CST, #12058) antibodies were purchased from Cell Signaling Technology. Anti-GATA4 (ab84593), anti-SIRT6 (ab62739), and anti-H3 (ab1791) antibodies were purchased from Abcam. Anti-P300 (sc48343) and anti-GST (sc138) antibodies were purchased from Santa Cruz Biotechnology. Anti-H3K9ac (ABE18) antibody was obtained from EMD Millipore. Anti-flag M2 antibodies were purchased from Sigma-Aldrich.

**Luciferase reporter assay**

HEK293 cells were transfected with a pGL3-Bcl-2-promoter reporter, the indicated expression plasmids and control plasmid pCMXβgal. Luciferase activities were measured using a luciferase reporter assay kit (Promega, #E1501) according to the manufacturer's instructions. The luciferase values were normalized according to β-gal activity.

**RNA preparation and Real-Time qPCR**

Total RNA was extracted from cells or mouse tissues using Trizol reagent RNAiso Plus (Takara) following the phenol-chloroform extraction method. cDNA library was synthesized from the purified RNA by Prime Script RT Master Mix (Takara) according to the method: 37 °C, 40 min; 85 °C, 5s. Gene expression was analyzed by CFX Connected™ Real-Time PCR Detection System (Bio-Rad) using a system of SYBR Ex Taq Premixes (Takara). Each experiment was performed at least in triplicate. Values were standardized to 18srRNA. The primers for qPCR are listed (Supplementary Table S1).

**Chromatin immunoprecipitation (ChIP)**

ChIP assay was performed as previously (1). Briefly, cells were cross-linked and lysed with lysis buffer (50mM Tris·HCl pH 8.0, 2mM EDTA, 15 mM NaCl, 1% SDS, 0.5% deoxycholate, protease inhibitor cocktail, and 1mM PMSF). After sonication and centrifugation, the supernatant was collected and precleared with protein A/G Sepharose. The precleared samples were incubated overnight with the relative antibodies, like anti-H3K9ac, anti-SIRT6, anti-GATA4 antibody (2 μg/sample) or appropriate IgGs as a control. The binding DNA fragments were precipitated by supplemented with 50% suspension of protein A/G Sepharose (Invitrogen). After washed sequentially with a series of buffers, the beads were heated at 65 °C to reverse the cross-link. DNA fragments were purified and analyzed by qPCR. The primers used in this study are listed (Supplementary Table S1).

**TUNEL staining assay**

Doxorubicin-induced cardiomyocyte apoptosis was evaluated by the Dead End™ Fluorometric TUNEL System (Promega, G3250) according to manufacturer's instructions. Five views were captured randomly under the microscope to calculate the TUNEL-positive staining rate for each slice.

**Fluorescence-activated cell sorting (FACS) assay**

H9C2 cells were treated with 1μM DOX for 12 hours, and then stained with FITC-Annexin V (Beyotime, C1062L) and 7-AAD (BD Pharmingen, 559925) for live/dead cell discrimination. Flow cytometry analyses were performed using an LSR-II flow cytometer and analyzed with FlowJo x7.5.

**Statistical Analysis**

All values are expressed as the mean ± SEM. Statistical differences among groups were determined using either Student's t test or two-way ANOVA using Graph-Pad Prism Software. Survival curve data were analyzed using log-rank (Mantel-Cox) test.

### Figure Legend

**Figure 1. SIRT6 enhances GATA4 transcription activity.**

(A) Quantitative PCR showing the mRNA levels of *Bcl-2*, *Bcl-xL*, *Sirt6*, and *Gata4* in heart tissues from Sirt6$^{flox/flox}$ and Sirt6$^{flox/flox}$; Myh6-Cre mice injected with tamoxifen. n=3/group. *P < 0.05, ***P < 0.001. 'ns' indicates no significance. (B) H9C2 cells were treated with 1 μM DOX for the indicated time points (0, 3, 6, 9hr). The mRNA levels of *Bcl-2*, *Sirt6*, and *Gata4* were determined by qPCR. ***P < 0.001, 'ns' indicates no significance. (C) H9C2 cells with si-NC, or si-*Sirt6* siRNAs were employed to DOX exposure for 4hr. The mRNA levels of *Bcl-2*, *Sirt6*, and *Gata4* were determined by qPCR. ***P < 0.001, 'ns' indicates no significance. (D) Quantitative PCR showing the mRNA levels of *Bcl-2*, *Bcl-xL*, and *Sirt6* in wild-type (WT) and *Sirt6* transgenic (*Sirt6*-Tg) mice after intraperitoneal injection of 10 mg/kg DOX twice in one week. n=5/group, *P < 0.05, **P < 0.01, ***P < 0.001. 'ns' indicates no significance. (E) Schematic of GATA consensus motif on *Bcl-2* promoter regions (Upper). P1 and P2 indicate -266 GATA motif and -1025 GATA site, respectively. TSS represents transcription start site of *Bcl-2* gene. ChIP analysis was performed using anti-SIRT6 antibody and IgG (Lower). The enrichment of SIRT6 on *Bcl-2* promoter region was analysis by qPCR with primers targeting P1 and P2 motif. *P < 0.05, **P < 0.01, ***P < 0.001. (F) ChIP analysis was performed using anti-GATA4 antibody in HEK293 cells with or without Flag-SIRT6 expression. The enrichment of GATA4 on *Bcl-2* promoter region was analysis by qPCR with primers targeting P1 motif. *P < 0.05, **P < 0.01, ***P < 0.001.

**Figure 2. SIRT6 physically interacts with GATA4**

(A) Co-immunoprecipitation (Co-IP) using anti-HA agarose was performed from HEK293 cells transfected with indicated constructs. The level of Flag-GATA4 was detected by immunoblotting in purified pool of HA-SIRT6. (B) Co-IP using anti-Flag agarose was performed from HEK293 cells transfected with indicated plasmids. HA-SIRT6 was tested by immunoblotting in Flag-GATA4 precipitates. (C) The plasmid of Flag-GATA4 was co-transfected with the truncated forms of HA-SIRT6 separately in HEK293 cells. Co-IP with anti-HA agarose was performed and Flag-GATA4 was determined by immunoblotting. ΔC indicates C-terminal deletion of SIRT6; ΔN is short of N-terminal deletion. (D) Purified *E. coli*-expressed His-GATA4 was incubated with GST-fused different fragments of SIRT6.

After GST pull-down experiment, the co-precipitated GATA4 was detected by immunoblotting. Coomassie blue staining indicates the loading level of GST-SIRT6. (E-F) The endogenous interaction of GATA4 and SIRT6 was evaluated by immunoprecipitation and immunostaining with anti-GATA4 and anti-SIRT6 antibodies. The immunoblots and representative photos of immunofluorescence staining were showed in Figure E and F, respectively. Scale bar, 50 µm. (G) Diagrams of Human GATA4 containing two Zn-finger DNA binding domains. (H) Domain-based truncations of HA-GATA4 were generated and co-expressed with Flag-SIRT6 in HEK293 cells. HA-GATA4 protein was precipitated and Flag-SIRT6 was analyzed by immunoblotting.

**Figure 3. SIRT6 enhances GATA4 acetylation independently of its deacylase activity.** (A) Flag-GATA4 co-expressed with nuclear sirtuins (SIRT1, SIRT2, SIRT6 and SIRT7) individually in HEK293 cells, and then was purified by anti-flag M2 beads. The acetylation level was determined with anti-acety-lysine (AcK) antibody. (B) The truncated constructs of HA-GATA4 were co-transfected with the plasmid of Flag-SIRT6 or empty vector in HEK293 cells. Immunoblotting was performed with the indicated antibodies. (C) HEK293 cells co-expressing Flag-GATA4 with HA-SIRT6 were treated with 10 µM NAM for 4hr. The acetylation level of GATA4 was examined using anti-AcK antibody after immunoprecipitation. (D) HA-GATA4 co-expressed with different forms of Flag-SIRT6 (WT, enzyme-dead mutation D63Y, and D116N) separately in HEK293 cells, and then was precipitated by anti-HA beads. The acetylation level was determined with anti-AcK antibody. The blot of Flag-SIRT6 in IPs indicated the association of GATA4 and SIRT6. (E) Real-time PCR analysis of mRNA levels of *Bcl-2* in WT and *Sirt6* null MEFs re-expressing Flag-SIRT6, or enzyme-dead mutation. *$P < 0.05$; **$P < 0.01$. 'ns' indicates no significance. 18srRNA was used as a reference gene. (F) Representative flow plots showing apoptotic H9C2 cells over-expressing the indicated proteins treated with 1µM DOX for 12 hours following FITC-Annexin V/7-AAD staining.

**Figure 4. SIRT6 recruits Tip60 for GATA4 acetylation**

(A) HEK293 cells ectopically expressing Flag-GATA4 were treated with various reagents to inhibit the relative HATs, like Cpth2, Mg149, and C646. The change on acetylation level of GATA4 was tested after immunoprecipitation. (B) HA-GATA4 co-expressed with Flag-Tip60 or empty vector in si-NC or si-*SIRT6* HEK293 cells. HA-GATA4 proteins were purified with anti-HA beads. The acetylation status of GATA4 was determined by immunoblotting with anti-AcK antibody. (C) Flag-GATA4 co-expressed with HA-SIRT6 or empty vector in HEK293 cells, which were pre-treated with si-NC or si-*Tip60*. The change of acetylation level of GATA4 was detected. (D) HEK293 cells were co-transfected Flag-Tip60 with HA-SIRT6, or empty vector. Flag-Tip60 proteins were purified. HA-SIRT6 protein and Tip60 acetylation level were examined by anti-HA and anti-AcK antibodies, respectively. (E) Immunoblotting showing the levels of endogenous GATA4 and SIRT6 in Flag-Tip60 IPs. H-C represents heavy chains of IgG. (F) Flag-Tip60 co-expressed with or without HA-GATA4 in wild-type (WT) or *SIRT6* KO HEK293 cells. Co-immunoprecipitation of HA-GATA4 was performed and the level of Flag-Tip60 was checked. (G) Alignment of protein sequence of human GATA4 and its orthologues in mouse, rat, and *Xenopus*. All lysine residues containing p300-targeted sites were highlighted. (H) The constructs of wild-type and site-mutated (K320/322R and K328/330R) HA-GATA4 were applied for co-transfection with Flag-SIRT6 plasmid. The acetylation levels of GATA4 were determined. (I) K328/330R mutation of GATA4 co-expressing with Flag-Tip60 or not were purified. The acetylation levels of GATA4 were determined by using anti-AcK antibody.

**Figure 5. Acetylation is critical for GATA4 preventing DOX-induced cardiotoxicity.** (A) HEK293 cells ectopically expressing HA-GATA4 and Flag-SIRT6 were subjected to doxorubicin treatment at different dosages for 6hr. Immunoblotting was performed to test the protein level of Flag-SIRT6 and acetylation level of GATA4 in the precipitated pool of HA-GATA4. (B) H9C2 cells stably expressing GATA4 WT, K328/330R (KR), K328/330Q (KQ) were employed to DOX-induced cardiotoxicity. The mRNA transcripts of *Bcl-2* were evaluated by qPCR. n = 3, *P < 0.05; **P < 0.01. 'ns' represents no significance. 18srRNA was used as a reference gene. (C) ChIP analysis was performed for these stably-transfected H9C2 cells by using anti-GATA4 antibody. The enrichment of GATA4 on *Bcl-2* promoter region was measured by qPCR. n ＞ 3, ***P < 0.001. (D-E) Stably transfected H9C2 cells were applied to DOX exposure for 24hr and then stained with crystal violet. The views of staining cells were randomly captured (D). Cell area was measured and quantified (E). Scale bar, 15 μm. n=8, **P < 0.01; ***P < 0.001, 'ns' represents no significance. (F) After treatment with DOX for 24hr, H9C2 cells were employed for TUNEL staining. The views were randomly captured, and the positive staining cells were calculated and analyzed. n＞ 6, ***P < 0.001, 'ns' indicates no significance. (G) H9C2 cells were treated with the indicated concentrations of DOX for 1hr and further grew in DOX-free medium for 1 week. Cell colonies in plates (n= 3/group) were stained with crystal violet. The colony numbers were calculated and statically analyzed as showed in the right table.

**Figure 6. *Sirt6* transgenic mice exhibits more resistant to DOX-induced cardiomyopathy.**

(A) Kaplan–Meier survival curves of wild-type (WT) and *Sirt6* transgenic (*Sirt6*-Tg) mice after intraperitoneal injection of 10 mg/kg DOX twice in one week. n=8, ***P < 0.001. Log-rank test. Arrows indicate DOX injection. (B) Body weight of mice at day7 after the last injection of DOX. n = 4 per group, *P < 0.05, 'ns' indicates no significance. (C) The ratio of heart weight to body weight (HW/BW) in WT and *Sirt6*-Tg mice were shown (n = 5 per group). *P < 0.05, 'ns' represents no significance. (D) The representative H&E staining of heart sections from WT and *Sirt6*-Tg mice at day7 after the last injection of DOX. Scale bar, 1mm. (E) Representative immunohistochemical images showing Gata4 proteins in heart sections from animals treated as in (A). Scale bar, 20 μm. (F) Relative quantification of cardiomyocyte area in heart sections from animals treated as in (A). ***P < 0.0001, 'ns' represents no significance. (G-H) The representative images and the relative quantification of collagen deposition with Masson staining in heart sections from animals treated as in (A). Scale bar, 100 μm. ***P < 0.0001, *P < 0.05. (I) Relative quantification of TUNEL-positive staining population in heart sections from animals treated as in (A). ***P < 0.0001.

**Figure 7. A schematic diagram of SIRT6-regulated GATA4 activity in protection against DOX-induced cardiotoxicity.**

Under normal condition (a), SIRT6 associates with GATA4 on C-tailed Zn-finger domain, and recruits Tip60 for GATA4 acetylation (red stars), promoting GATA4 transcriptional outcome and maintaining cardiomyocyte survival. When exposure to doxorubicin (DOX) (b), SIRT6 expression and the trimetric complex are disrupted, consequently blocking GATA4 transcription activity and triggering myocyte apoptosis.

### Supplementary Figure Legend

**Figure S1. SIRT6 enhances GATA4 chromatin binding affinity.**

(A) Volcano plot of comparing the gene/repeat expression profiles between wild-type and

*Sirt6* null MEFs. Plotted for each transcript are the negative log10 of the p value and the log2 of the fold change of gene expression of *Sirt6* null versus WT cells. FC, fold change. (B) Immunoblotting showing the protein levels of Gata4 and Sirt6 in wild-type (WT) and *Sirt6* depleted (KO) mouse hearts. GAPDH as a loading control. (C) H9C2 cells were transfected with siRNAs targeting *Sirt6*, or *Gata4* expression. The transcripts of *Bcl-2* and *Bcl-xL* were analyzed by qPCR. *P < 0.05; ***P < 0.001. 'ns' represents no significance. (D) HEK293 cells were transfected with the plasmids as indicated, and then the values of luciferase activities were measured and normalized to β-gal activities. **P < 0.01. (E) ChIP analysis was performed using anti-H3K9ac antibody in HEK293 cells with or without Flag-SIRT6 expression. The enrichment of H3K9ac on *Bcl-2* promoter region was analysis by qPCR with primers targeting P1 motif. *P < 0.05, **P < 0.01, ***P < 0.001. (F) Immunoblotting showing the levels of Flag-GATA4 in chromatin-bound fractions (P2) and total cellular extracts (TCE) from HEK293 cells with or without HA-SIRT6 over-expression. (G) Immunoblotting showing the levels of endogenous GATA4 in chromatin-bound fractions (P2) and total cellular extracts (TCE) from si-NC or si-*SIRT6* HEK293 cells.

**Figure S2. SIRT6 enhances GATA4 acetylation and attenuates DOX-induced myocyte apoptosis.**

(A) Immunoblotting showing the acetylation level of GATA4 after immunoprecipitation of HA-GATA4 from si-NC or si-*SIRT6* HEK293 cells. "#1" and "#2" represent two independent sequences of siRNAs targeting *SIRT6* transcript. (B) HEK293 cells transfected with the indicated plasmids were collected. After immunoprecipitation, the acetylation of GATA4 were determined by anti-AcK antibody. (C) Co-immunoprecipitation using anti-Flag agarose was performed from HEK293 cells expressing HA-GATA4 alone, or with Flag-SIRT6 wild-type or H133Y mutation. The level of HA-GATA4 was tested in Flag-SIRT6 precipitates by immunoblotting. (D) Representative flow plots showing apoptotic H9C2 cells over-expressing the indicated proteins treated with 1μM DOX for 12 hours following FITC-Annexin V/7-AAD staining.

**Figure S3. Tip60 involves SIRT6-enhanced acetylation of GATA4.**

(A) HEK293 cells expressing HA-GATA4 alone or with Flag-Tip60 were treated with Mg149 (0, 10 or 20 μM) for 6hr. After extraction and immunoprecipitation, the acetylation levels of GATA4 were checked by immunoblotting. (B) HEK293 cells expressing HA-GATA4 were transfected with si-NC, si-*Tip60*, or si-*p300* siRNAs. The acetylation levels of GATA4 were tested by immunoblotting after immunoprecipitation. (C) Co-immunoprecipitation with anti-SIRT6 antibody was performed in HEK293 cells expressing Flag-Tip60 alone, or with Flag-SIRT6 wild-type or site-mutations. The level of Flag-Tip60 was tested in the anti-SIRT6 precipitates.

**Figure S4. GATA4 KR mutation dampens its protective function against DOX.**

(A) Immunoblotting identifying the expression of different forms of GATA4 in stably-transfected H9C2 cells. "Ve" indicates empty vector transfection. WT, KR, and KQ represent GATA4 wild-type, K328/330R mutation, and K328/330Q mutation, separately. (B) Immunoblotting showing the level of GATA4 in chromatin-bound fractions (Chr) and whole cellular lysates (WCL) from stably-transfected H9C2 cells. (C) The representative images showing the TUNEL-staining in stably-transfected H9C2 cells after DOX or DMSO treatment. The relative quantification data was shown in Figure 5F. Scale bar, 50 μm.

**Figure S5. *Sirt6* transgenic mice exhibits better tolerance to DOX exposure.**

(A) Representative images showing the whole heart sizes harvested from wild-type (WT) and *Sirt6* transgenic (*Sirt6*-Tg) mice with or without intraperitoneal injection of DOX. (B) The relative quantification showing Gata4 proteins in heart sections in Figure 6E. *$P < 0.05$, 'ns' indicates no significance. (C) The representative images showing H&E staining in heart sections from animals treated as in Figure 6A. Relative quantification of cardiomyocyte area was shown in Figure 6F. Scale bar, 50 μm. (D) The representative images showing TUNEL staining in heart sections from animals treated as in Figure 6A. Relative quantification of TUNEL-positive staining population was shown in Figure 6I. Scale bar, 20 μm.


1. Qian, M., Liu, Z., Peng, L., Tang, X., Meng, F., Ao, Y., Zhou, M., Wang, M., Cao, X., Qin, B. *et al.* (2018) Boosting ATM activity alleviates aging and extends lifespan in a mouse model of progeria. *eLife*, **7**.
2. Qian, M.X., Pang, Y., Liu, C.H., Haratake, K., Du, B.Y., Ji, D.Y., Wang, G.F., Zhu, Q.Q., Song, W., Yu, Y. *et al.* (2013) Acetylation-mediated proteasomal degradation of core histones during DNA repair and spermatogenesis. *Cell*, **153**, 1012-1024.
3. Kobayashi, S., Lackey, T., Huang, Y., Bisping, E., Pu, W.T., Boxer, L.M. and Liang, Q. (2006) Transcription factor gata4 regulates cardiac BCL2 gene expression in vitro and in vivo. *FASEB journal : official publication of the Federation of American Societies for Experimental Biology*, **20**, 800-802.
